# Microfluidic guillotine reveals multiple timescales and mechanical modes of wound response in *Stentor coeruleus*

**DOI:** 10.1101/2020.11.12.379123

**Authors:** Kevin S. Zhang, Lucas R. Blauch, Wesley Huang, Wallace F. Marshall, Sindy K. Y. Tang

## Abstract

**Background:** Wound healing is one of the defining features of life and is seen not only in tissues but also within individual cells. Understanding wound response at the single-cell level is critical for determining fundamental cellular functions needed for cell repair and survival. This understanding could also enable the engineering of single-cell wound repair strategies in emerging synthetic cell research. One approach is to examine and adapt self-repair mechanisms from a living system that already demonstrates robust capacity to heal from large wounds. Towards this end, *Stentor coeruleus*, a single-celled free-living ciliate protozoan, is a unique model because of its robust wound healing capacity. This capacity allows one to perturb the wounding conditions and measure their effect on the repair process without immediately causing cell death, thereby providing a robust platform for probing the self-repair mechanism.

**Results:** Here we used a microfluidic guillotine and a fluorescence-based assay to probe the timescales and mechanisms of wound repair in *Stentor*. We found that *Stentor* requires ∼100 – 1000 s to close bisection wounds, depending on the severity of the wound. This corresponds to a healing rate of ∼8 – 80 μm^2^/s, faster than most other single cells reported in the literature. Further, we observed and characterized three distinct mechanical modes of wound repair in *Stentor*: contraction, cytoplasm retrieval, and twisting/pulling. Using chemical perturbations, active cilia were found to be important for only the twisting/pulling mode. Contraction of myonemes, a major contractile fiber in *Stentor*, was surprisingly not important for the contraction mode and was of low importance for the others.

**Conclusions:** While events local to the wound site have been the focus of many single-cell wound repair studies, our results suggest that large-scale mechanical behaviors may be of greater importance to single-cell wound repair than previously thought. The work here advances our understanding of the wound response in *Stentor*, and will lay the foundation for further investigations into the underlying components and molecular mechanisms involved.

## 1. Background

Wound repair is a fundamental property of life. It is an essential biological process for maintaining homeostasis and, ultimately, for survival. While wound repair is known to occur at the tissue level, there is increasing recognition that single cells also have a wound response. Single-cell wound repair has been reported in fungi, amoebae, budding yeast, and also the Metazoa [1], [2], [3]. Understanding wound repair at the single-cell level is thus critical for elucidating important cellular functions for homeostasis and survival across the kingdoms of life, and also ultimately for developing therapeutic approaches for wound-induced diseases [1].

From a different angle, the ability to engineer single-cell wound repair strategies can find use in emerging synthetic cell research. Recent years have witnessed an explosion in research on building synthetic cells [4], [5], [6], not only as an experimental method to study the origins and the rules of life, but also as a new approach to biochemical engineering, in which molecules (e.g., enzymes) are encapsulated in membranes to increase local concentrations and regulate substrate/product exchange, leading to significant increases in the yield and specificity of reactions [7], [8]. Nevertheless, current synthetic cell research has largely neglected one of the most fundamental properties of living matter – the ability to self-repair following damages. If such self-repairing capability can be introduced in synthetic cells, it could open new realms of biochemical engineering by allowing the synthetic cell systems to operate robustly under the potentially harsh environment of industrial processes.

One approach to attaining self-repairing synthetic cells is to adapt self-repair mechanisms from a living system that already demonstrates robust capacity to heal from large mechanical wounds within a single cell, and build analogs of these mechanisms inside synthetic cells. One of such systems is *Stentor coeruleus*, a single-celled free-living ciliate protozoan. Our rationales for studying wound healing in *Stentor* are: 1) Its wound healing capacity is more robust than most other cells (see details in Table 1). *Stentor* possesses a highly polyploid macronucleus such that even small cell fragments, as small as 1/27^th^ of original cell size, can contain enough genomic copies to survive and regenerate in 24 hours [9, 10]. This unique wound healing property allows us to perturb the wounding conditions and measure their effect on the repair process without immediately causing cell death, thereby providing a robust platform for probing the self-repair mechanism. 2) The ability to perform high-throughput gene knockdown and wounding experiments. We recently sequenced the *Stentor* genome [11] and developed tools for molecular manipulation of *Stentor* gene expression [12], thus paving the way to a molecular understanding of *Stentor* wound repair.

**Table 1:** Summary of wound repair timescales in single cells

| Cell type | Wound diameter (μm) | Cell diameter (μm) | Ratio of wound diameter to cell diameter | Healing time (s) | Approx. healing rate (μm<sup>2</sup>/s) |
| --- | --- | --- | --- | --- | --- |
| LHCN skeletal muscle (human)<sup>[43]</sup> | 1 | ~40 | 0.03 | 70 | 0.01 |
| HeLa[44] | 2-3 | 30 | 0.07 | 200-300 | 0.01-0.03 |
| 3T3 fibroblast[45] | 2 | 18 | 0.11 | 60-120 | 0.02-0.05 |
| Dictyostelium[46] | 0.5-2 | 15 | 0.03 | 5-10 | 0.02-0.6 |
| Drosophila embryo[47] | 12-20 | 150-500 | 0.08 | 100-200 | 0.6-3.1 |
| Xenopus oocyte[48] | ~200 | 1200 | 0.17 | 600 | 52 |
| Sea urchin egg[49] [50] [51] | 10 – 50 | 80 | 0.125 – 0.625 | 3 – 15 | 31 – 130 |
| ***Stentor coeruleus*** | **~100** | **~200**<br>**(for a cell fragment approx. half the original cell size)** | **0.5** | **100 – 1000** | **7.9 – 79** |

Previous studies of single-cell wound response in organisms such as *Xenopus* oocytes and *Drosophila* embryos have indicated that the healing of plasma membrane wounds involves active cellular processes. At least two steps are involved when the plasma membrane is disrupted: 1) the influx of calcium ions through the membrane opening triggers the sealing of the plasma membrane via active trafficking of internal membranes to the wound site, and 2) active remodeling of the cytoskeleton [1], [13], [14], [15]. In some organisms, the latter process involves actin accumulation around the wounds and the formation of a contractile purse string, as well as the recruitment of Rho to the plasma membrane and the subsequent active actin assembly at sites of Rho activation [16], [17], [18].

Currently, it is unknown whether *Stentor* also employs similar processes to heal wounds. As *Stentor* is a ciliate with cellular structures that are distinct from the organisms that have been studied previously for single-cell repair, it is possible that *Stentor* utilizes different mechanisms to repair wounds. Structurally, *Stentor* is covered in cilia for locomotion. It consists of an oral apparatus with a dense membranellar band of cilia around the anterior of the cell for feeding. It possesses a holdfast, an anchoring structure, at its posterior. The cell cortex is defined by the oral apparatus and the holdfast, together with ciliated stripes that run parallel to the long axis of the organism. Electron micrographs of *Stentor* revealed that the ciliated stripes consist of rows of cortical fibers comprising two primary types of filaments, the KM fibers and myonemes [19]. KM fibers are made up of bundles of microtubules, and are attached to the cell membrane via basal bodies. Myonemes, located immediately under the KM fibers, are contractile fibers responsible for the rapid contraction of the cell from an extended trumpet shape to a sphere at rates up to 10 – 20 cm/s [20], [21]. Immunostaining of *Stentor* showed the localization of a protein that is immunologically related to centrin/caltractin, a class of EF-hand calcium binding proteins that form contractile filaments in a variety of organisms, in myonemes [22]. Further details on the anatomy of *Stentor* can be found in previous work [23].

In this paper, we aim to quantify the healing time and characterize the wound response in *Stentor* inflicted with mechanical wounds from a microfluidic “guillotine”. Previously, we have developed a microfluidic guillotine for high throughput wounding and bisection of *Stentor* cells in a continuous flow manner [24]. Changing the applied flow velocity changes the local cutting dynamics, and leads to two regimes of cell bisection: Regime 1 at low viscous stress where cells are cleanly cut with small membrane ruptures and high viability (∼97%), and Regime 2 at high viscous stress where cells are torn with extended membrane ruptures and decreased viability (∼60% - 80%). While laser ablation allows more exact control of wound size and location compared with our method, it requires immobilizing *Stentor* which has been challenging since *Stentor* tends to swim away or contract upon laser excitation. The continuous flow design of our guillotine circumvents this issue while also allowing us to probe the wound healing characteristics of a large number of cells from seconds to hours after the cells are wounded. Combined with a fluorescence-based assay to detect the presence or absence of a wound, we show that *Stentor* takes ∼100 – 1000 seconds to heal their wounds, using three mechanical modes of wound response that have not been reported in other cell types. The prevalence of these three mechanical modes in *Stentor* suggests that mechanical behaviors such as cellular force generation and motility could play a more important role in single-cell wound repair than previously thought. This work is expected to lay the foundation for further investigations on the molecular mechanisms of wound repair in *Stentor*.

## 2. Results

### 2.1. Design of the wound repair assay using a parallelized microfluidic guillotine device and Sytox Green staining

To study the wound repair process in *Stentor coeruleus*, we designed our microfluidic device with 8 parallelized guillotine channels arranged in a radial geometry converging at a single outlet. This design reduced the distance each cell had to travel and the corresponding lag time prior to assaying the cells. Figure 1A shows a schematic diagram of our parallelized guillotine device. The cell suspension was injected via a cell inlet (constant flow rate of 8 and 36 mL/hr corresponding to an average velocity of 1.4 and 6.3 cm/s per guillotine for Regime 1 and 2, respectively). A flushing inlet was used to inject cell media (Pasteurized Spring Water, PSW) only and was connected directly to the outlet to control the lag time prior to assaying the cells (see details in Section 5.3).

**Figure 1.**
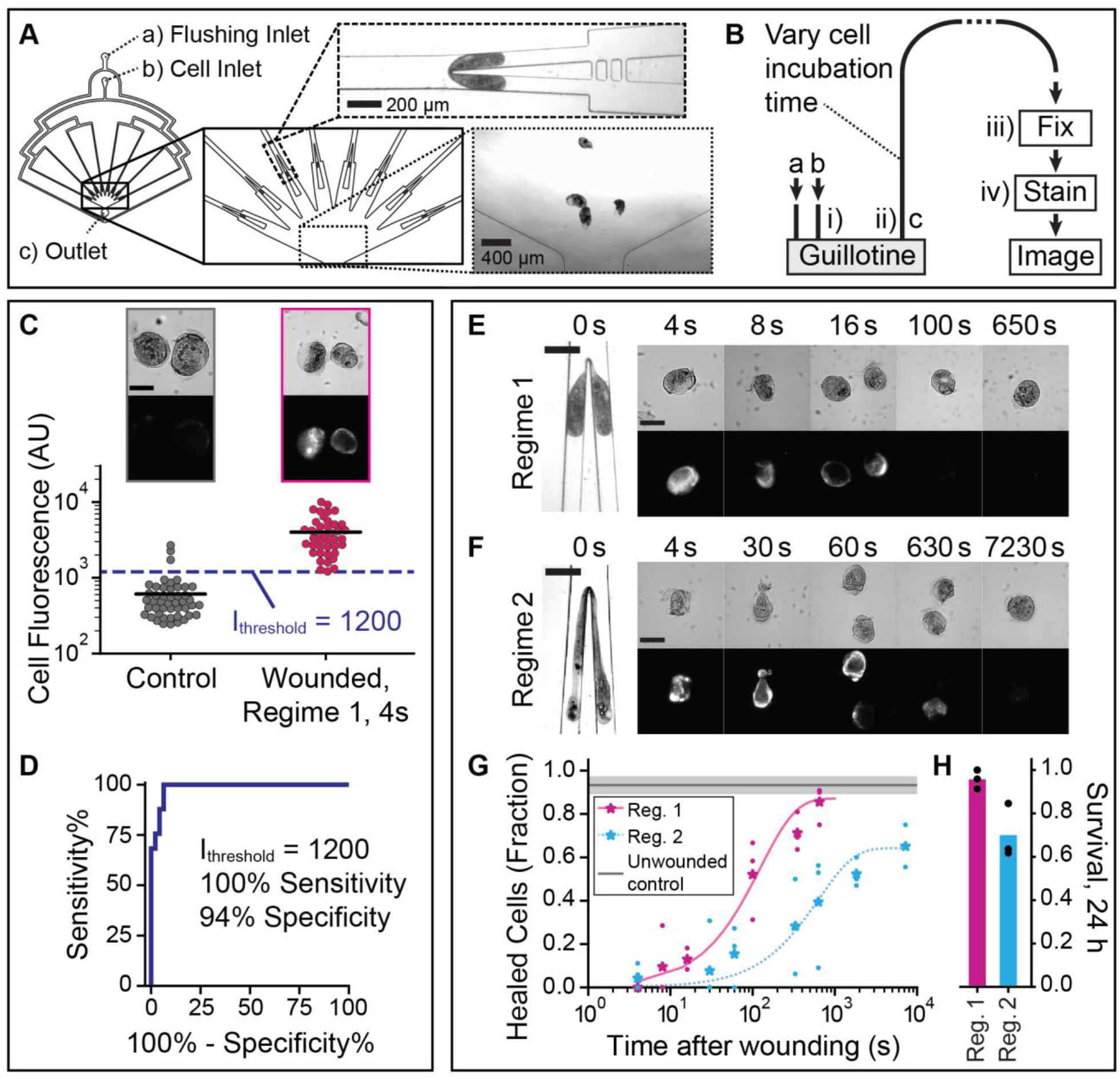
Wound repair assay design and characterization of wound repair in *Stentor coeruleus*. Parallelized guillotine device, with two inlets, **a** and **b**, used for flushing media and injecting cells, respectively, and one outlet **c**. To parallelize cell cutting, a radial array of 8 guillotines converged in a well that led to the outlet **c**. Schematic diagram of the wound repair assay. **a, b**, and **c** refer to the inlets/outlets. i-iv) are steps in the assay (See Sections 2.1 and 5.3-4). **C)** Cell fluorescence for unwounded cells (controls) (N=46 cells) and wounded cells (Regime 1, 4 s post-wounding) (N=41 cells), using the wound repair assay. Data was combined from 3 biological replicates per experimental group, with mean lines shown. Representative brightfield and fluorescence images shown. The dashed line at cell fluorescence=1200 AU indicates the intensity threshold used to distinguish wounded from healed cells, I_threshold_. **D)** ROC (Receiver Operating Characteristic) curve for the wound repair assay. Representative images of cells wounded in **E)** Regime 1 and **F)** Regime 2: brightfield (top panels) and corresponding fluorescence images (bottom panels) of wounded cells, fixed and stained with Sytox Green at different time points post-wounding. All fluorescence images were scaled equally (100-10000 AU). Scale bars in **C), E)**, and **F)** are 200 μm. **G)** Fraction of cells healed, using the wound repair assay. Data shown as mean (stars) of N≥3 biological replicates (dots) and fit to a one-phase exponential function using mean datapoints in addition to the 24-hour (86400 s) survival rate, to aid extrapolation. The control line shows the mean fraction of unwounded control cells below I_threshold_, and the shaded region indicates standard deviation (SD) (3 biological replicates). Total N=31-65 cells per experimental condition. **H)** Survival rate of cells wounded in Regimes 1 and 2, 24 hours after wounding, shown as mean (bar) of 3 biological replicates (dots) (N=22-34 cells per replicate).

Our wound repair assay involved 4 steps (Figure 1B): i) injecting the cells into the guillotine and wounding them, ii) collecting the wounded cells via an outlet tubing, iii) fixing the cells, and iv) staining the fixed cells with Sytox Green followed by fluorescence imaging to determine if the wound was open or closed. In the time t_post-wound_ between cell wounding at the guillotine and when the cells were fixed (i.e. the duration of step ii), wounded cells had the opportunity to repair their wounds. By varying t_post-wound_ (see details in Section 5.3), we expected to assess the completeness of wound repair as a function of time. In this work we varied t_post-wound_ from 4 seconds up to 150 minutes. The lower limit was set by the minimum tubing length that could be used practically in our experiment and the highest flow rate we could apply without further wounding the cells. Our results (Figure 1E – H) indicated that this timing resolution was sufficient to capture the wound repair dynamics, which occurred over hundreds of seconds.

As no membrane dye to-date has worked for staining the plasma membrane of *Stentor*, we developed an indirect assay to estimate the completion of wound repair by measuring the fluorescence intensity of a cell-impermeable dye Sytox Green inside a wounded cell. Sytox Green is commonly used to stain nucleic acids in dead cells, which have permeabilized membranes [25], [26], [27]. We found that on fixed *Stentor* cells, Sytox Green stained wounded cells, likely due to the presence of nucleic acids (RNA) in the cytosol of *Stentor*, but did not stain unwounded control cells. Figure 1C shows that the mean fluorescence intensity of wounded cells (4 seconds after their wounding) was approximately 10 times higher than that of the unwounded control cells. Sytox also sometimes brightly stained the macronucleus (DNA) of highly wounded cells. Although this indirect approach cannot give an absolute measurement of the wound size, the presence or absence of fluorescence could still indicate the presence or absence of a wound. It can thus be used for the binary classification of cells that were unwounded or completely repaired, *versus* those that were still wounded and thus permeable to Sytox Green at the time of the fixing step. Using a threshold intensity of 1200 (arbitrary units, AU), we were able to distinguish wounded cells from unwounded ones with 100% sensitivity and 94% specificity (Figure 1D).

### 2.2. Cells wounded in Regime 2 took longer to heal than those wounded in Regime 1

Figures 1E – F show representative images of cells being cut at the guillotine, along with fluorescence images of Sytox staining in representative cell fragments at different t_post-wound_ in Regimes 1 and 2 respectively. To quantify the wound healing results, we used the mean fluorescence intensity threshold I_threshold_ of 1200 (arbitrary units) for classifying healed cells vs. wounded cells (Figure 1C), and plotted the fraction of cells healed as a function of t_post-wound_. Figure 1G shows that 50% of the cells cut in Regimes 1 and 2 healed in about 100 seconds and 1000 seconds, respectively. We report the mean percentage of cells healed at each t_post-wound_ from at least 3 independent biological replicates. The individual fluorescence intensities of Sytox stained cells at different time points are shown in Additional File 1: Figure S2. In each regime, the maximum percentage of cells healed as measured using the wound repair assay (Figure 1G) plateaued to a value consistent with the survival rate of cells measured at 24 hours after wounding (Figure 1H).

Immunofluorescence images of acetylated tubulin on cell fragments revealed more details on the damage to the cytoskeleton of the wounded cells (Figures 2B – C) compared with unwounded cells (Figure 2A). As the KM fibers, which are microtubule structures arranged in cortical rows, are positioned directly under the cell membrane (within ∼1 μm [19]), the absence of immunostaining of acetylated tubulin (red arrowheads) could indicate a wound to both the membrane and the cortex, though it has to be verified using Sytox staining. The discontinuities or misalignment in the cortical rows of microtubules indicate that the membrane wound could be closed but the cortical rows had not been reorganized to restore the normal orientation.

**Figure 2.**
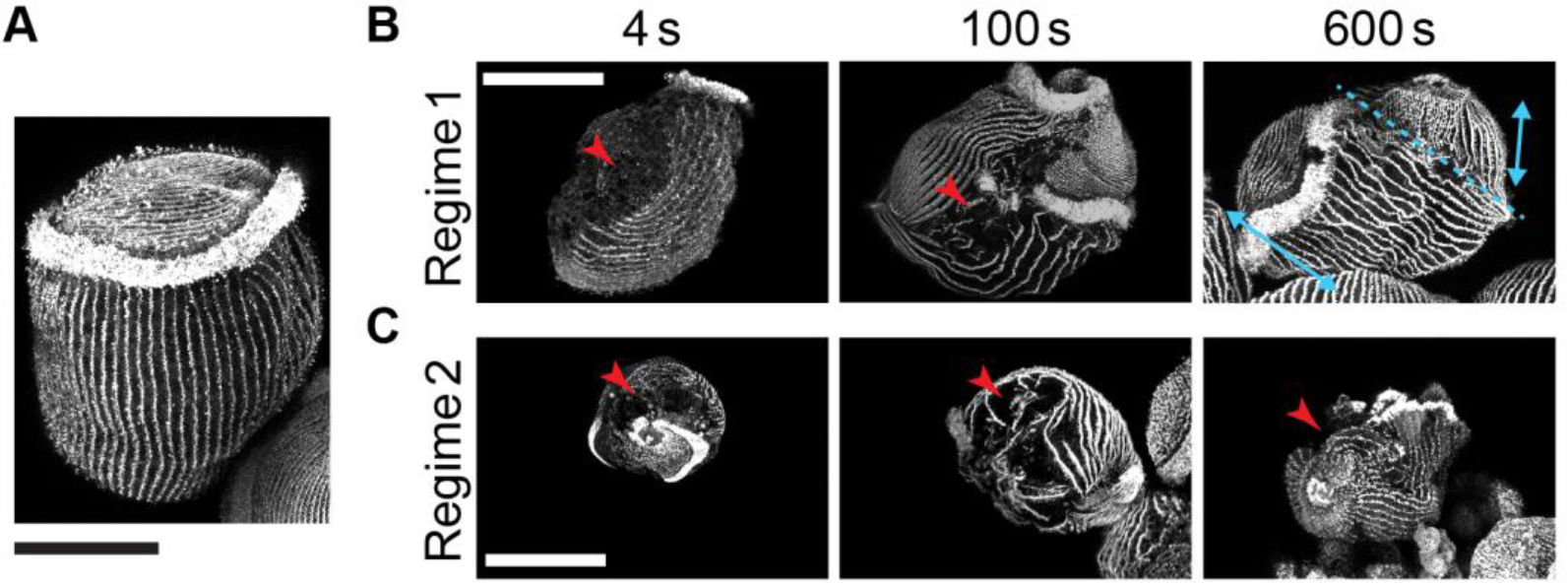
Immunostaining of acetylated tubulin. **A)** Representative immunostaining image of KM fibers in an unwounded cell. **B)** Representative immunostaining images of KM fibers in cells wounded in Regime 1. Red arrowheads point to the possible location of a wound. The dashed blue line in the healed cell at 10 min indicates the possible location of folding, as seen by the discontinuity in the KM fibers on either side of this line (blue arrows). Representative immunostaining images of KM fibers in cells wounded in Regime 2. All fluorescence images were acquired on the inverted confocal microscope. All scale bars 100 μm.

Overall, the tubulin staining images showed similar trends as the Sytox staining images. For example, cells cut in Regime 1 did not have any observable gaps in their cortical rows by t_post-wound_ =10 minutes, consistent with the lack of fluorescence in the Sytox staining at that time point. The wounds at earlier time points appeared to be localized to one distinct location. On the other hand, cells cut in Regime 2 had multiple gaps in their cortical rows at t_post-wound_ = 4 and 100 seconds, suggesting that there could be multiple wound sites. In addition, the misalignment in the cortical rows was more severe compared with those in Regime 1. The oral apparatus of the cell, which was stained very brightly by the immunofluorescence of acetylated tubulin, was often observed in multiple pieces over the cell or was completely absent (Additional File 1: Figure S1). Additional examples of wounded cells and tubulin staining are shown in Additional File 1: Figure S1.

### 2.3. Mechanical modes of wound response

Unlike most single-cell wound healing models studied previously, *Stentor* use their motile cilia to achieve a high degree of motility [23]. By observing cells after wounding, we identified a range of cell motions, which could be grouped into three mechanical modes of wound response (see Additional Files 2 – 5: Movies S1a – d).

#### Contraction

Contraction was observed in cells with a wound localized to one side of the cell. The cell contracted or folded around the wound site to reduce the wound size and eventually to close it (Figure 3A). Contraction often involved the entire cell, as evidenced by a decrease in cell length during the process. Due to the limited resolution of our imaging setup, we considered contraction complete when the wound diameter was reduced to ∼ 20 μm. Contraction typically took ∼100 – 250 seconds. When complete, the cells appeared folded. Additional File 1: Figure S3 details the analysis of the contraction mode.

**Figure 3.**
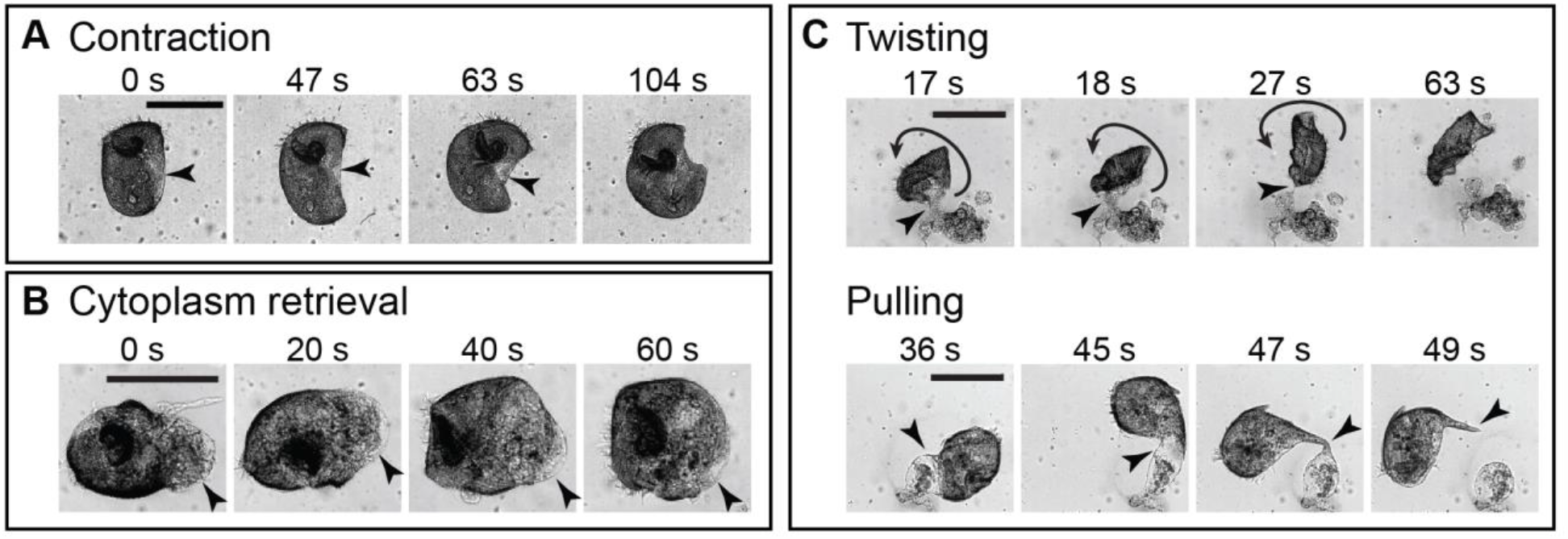
Mechanical modes of wound response. **A)** Representative images of a cell wounded in Regime 1 undergoing contraction. The arrows point to the location of the wound. **B)** Representative images of a cell wounded in Regime 2 undergoing cytoplasm retrieval. The arrows point to the retrieved cytoplasm that is part of the wound. **C)** Representative images of cells wounded in Regime 2 utilizing twisting/pulling. The short arrows point to the location of the wound, while the curved arrow indicates the direction of twisting. To better visualize the response in four panels, we did not include images at the beginning of cell twisting/pulling. All scale bars 200 μm.

The effects of the folding behavior observed during contraction could also be seen in the immunostaining of acetylated tubulin. The discontinuity in cortical rows (indicated by the blue arrows and dashed line in Figure 2B, t_post-wound_ = 10 min; additional images in Additional File 1: Figure S1), where the bottom set of cortical rows appeared perpendicular to the set at the top-right, is consistent with the cell folding onto itself to close the wound. We found that contraction occurred in more than 80% of the cells observed in Regime 1 (N=22/26 cells), as Regime 1 tended to generate single, localized wounds. While less common in Regime 2, contraction still occurred in ∼20% of cells observed (N=6/31 cells), only when the wound was localized to one side of the cell like in Regime 1. However, wounding was typically more severe in Regime 2 than in Regime 1, with multiple wounds observable to the eye and a larger amount of extruded cytoplasm, which may have prevented the contraction mode from occurring.

#### Cytoplasm retrieval

We also observed cells retrieving extruded cytoplasm into the cell (Figure 3B). The retrieval of cytoplasm appeared to follow a change in the shape of the cell. The retrieval process took ∼20 – 200 seconds. In Regime 1, cytoplasm retrieval occurred in about 50% of the cells observed (N=14/26 cells) and often occurred together with contraction (N=11/26 cells). Cytoplasm retrieval often occurred prior to or during the contraction healing mode. Cytoplasm retrieval occurred in about 35% of the cells observed in Regime 2 (N=11/31 cells). These cells typically did not retrieve all of the extruded cytoplasm like they did in Regime 1. Instead, some cytoplasm separated during the retrieval process and remained as debris.

#### Twisting/Pulling

We found that *Stentor* also used twisting/pulling motions to facilitate wound closure. For cells wounded in Regime 2 where part of their extruded cytoplasm became immobilized on the substrate, twisting and pulling motions were often observed to detach the cell from its lost cytoplasm (Figure 3C).

In the twisting mode, the less wounded part of the cell, which still contained beating cilia, twisted repeatedly and resulted in the supercoiling and eventual pinching of the wound site, thereby freeing the cell from the extruded cytoplasm. In the pulling mode, the more intact part of the cell was observed to swim and pull away from the extruded cytoplasm without twisting. The pulling motion formed a thin fiber at the point of detachment. At times, these thin fibers were seen to fold back over the rest of the cell.

Twisting/pulling healing modes typically completed in ∼20 – 100 seconds. We rarely observed twisting/pulling healing modes in cells wounded in Regime 1 (N=1/26 cells). In Regime 2, twisting/pulling healing modes occurred in about 60% of the cells (N=18/31 cells), and occasionally occurred with either contraction or cytoplasm retrieval. In cells that used the twisting/pulling mode, the pulling mode (N=15/19 cells) occurred more often than twisting (N=8/19 cells), with 4 cells utilizing both modes.

Among all the cells observed here, cells that had extruded cytoplasm always utilized cytoplasm retrieval, twisting/pulling, or both. Finally, we note that we did not observe any cell utilizing all three modes during the wound repair process. ∼5% of cells in Regime 1 (N=1/26 cells) and 10% of cells in Regime 2 (N=3/31 cells) were observed to not utilize any mechanical modes in their wound response.

### 2.4. Effect of chemical perturbations on wound response

To identify cellular components or factors contributing to the mechanical modes of wound response, we investigated the effects of two chemical agents, nickel chloride (NiCl_2_) and potassium iodide (KI), which were previously reported to inhibit cilia motion in *Tetrahymena* and *Paramecium* by inhibiting axonemal dynein [28], [29] and myoneme contraction in *Stentor* [23] respectively. NiSO_4_, another Ni^2+^ salt, was reported to have inhibitory effects on *Stentor* cilia [23].

In unwounded and untreated control cells, the beating cilia bundles on the membranellar band (Figure 4A) produced metachronal waves that appeared in the 2D autocorrelations as parallel lines (Figure 4A – C, Additional File 6: Movie S2a). These lines corresponded to a mean wave propagation speed of ∼1.09 ± 0.14 (standard deviation) mm/s and cilia beat frequency of ∼22.9 ± 3.7 Hz (N=6 cells). Upon wounding in Regime 2, the membranellar band of untreated control cells continued to beat in a coordinated fashion at similar speed and frequency as the unwounded case, with a mean wave propagation speed of ∼1.17 ± 0.32 mm/s and cilia beat frequency of ∼21.3 ± 7.0 Hz (N=6 cells) (Figure 4B – C, Additional File 7: Movie S2b).

**Figure 4.**
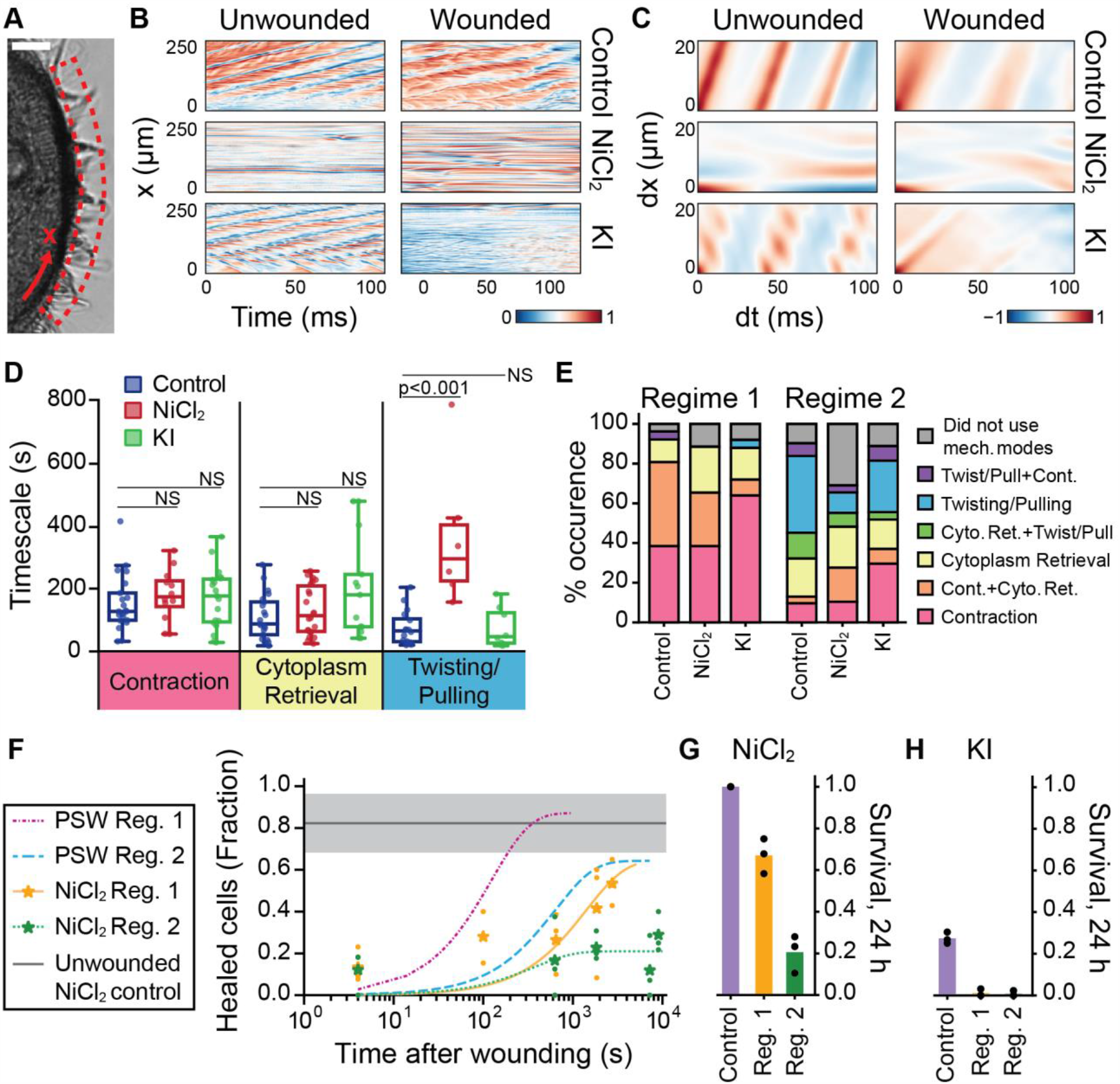
Effect of chemical perturbations on wound response. **A)** Region of interest and 1-D coordinate system defined along the membranellar band. Scale bar 20 μm. **B)** Kymographs and **C)** 2-D autocorrelations of the kymographs for cells under control (no treatment), NiCl_2_, or KI treatment, before and after wounding in Regime 2. Analysis of representative cells shown (N=6 per experimental group). **D)** Measured timescales of mechanical modes of wound response in untreated control, NiCl_2_-treated, and KI-treated cells, shown as box and whisker plots with a line at the median. Data in each treatment condition combined from Regime 1 and 2; individual cells shown as dots (Control: N=16-21 cells per mode; NiCl_2_: N=6-22 cells per mode; KI: N=8-19 cells per mode). Timescales were measured as described in Section 5.6. Significance values were obtained from multiple t-tests for treated cells compared to untreated cells, α=0.05. Only the NiCl_2_-treated twisting/pulling timescale was significantly different from that of untreated cells (p=0.0006). All other timescales were not significantly different (NS, p>0.05). **E)** Relative occurrences of mechanical modes of wound response across wounding regime and treatment condition (Total N=25-31 cells per experimental group). **F)** Fraction of NiCl_2_-treated cells healed, using the wound repair assay. Data shown as mean (stars) of N≥3 biological replicates (dots) and fit to a one-phase exponential function using mean datapoints in addition to the 24-hour (86400 s) survival rate, to aid extrapolation. Untreated (PSW) curves included for comparison. The control line shows the mean fraction of unwounded NiCl_2_-treated control cells below I_threshold_, and the shaded region indicates SD (3 biological replicates). Total N=26-59 cells per experimental condition. Survival rate of **G)** NiCl_2_-treated cells and **H)** KI-treated cells. Data for unwounded control cells, 24 hours after washing, and Regime 1 or 2 wounded cells, 24 hours after wounding, shown as mean (bar) of 3 biological replicates (dots) (N=15-47 cells per replicate).

First, we tested the effect of NiCl_2_ treatment. We verified that NiCl_2_ treatment suppressed cilia beating in unwounded cells (N=6 cells) and that the suppression persisted after wounding (N=6 cells), evidenced by a lack of motion in the kymograph and the absence of a clear wave pattern in the 2D autocorrelation (Figure 4B – C, Additional File 8 – 9: Movies S2c – d). Accordingly, the twisting/pulling mode of repair was less frequently observed in NiCl_2_-treated cells (∼20% in Regime 2, N=6/29 cells) compared with untreated cells (∼60% in Regime 2, N=18/31 cells) (Figure 4E). When a twisting/pulling mode occurred in a NiCl_2_-treated cell, it took significantly longer (∼3 times longer) for the wounded cell to detach itself from the extruded cytoplasm compared to untreated cells (p=0.0006) (Figure 4D). NiCl_2_ treatment had an insignificant effect on the other modes of wound response, contraction (p=0.84) and cytoplasm retrieval (p=0.84). These results indicated that cilia motion was primarily responsible for the twisting and pulling motions. Compared to ∼10% of Regime 2 untreated control cells, ∼30% of Regime 2 NiCl_2_-treated cells (N=9/29 cells) did not appear to utilize a mechanical mode of wound response. In Regime 2, while the wound healing time as assessed by the wound repair assay remained ∼1000 s (Figure 4F), the average NiCl_2_-treated cell survival rate was greatly reduced to 20% (Figure 4G), compared with 70% for untreated cells (Figure 1H). The survival of unwounded NiCl_2_-treated cells was not affected (100% survival) (Figure 4G). The average NiCl_2_-treated cell survival rate in Regime 1 was 67% compared with 96% for untreated cells (Figure 4G, Figure 1H), which was a lesser reduction in survival rate than seen in Regime 2. Surprisingly, in Regime 1 the wound healing time for NiCl_2_-treated cells, as assessed by the wound repair assay, increased significantly from ∼100 s to ∼1000 s (Figure 4F), which may have been due to off-target effects or toxicity of Ni^2+^. The individual fluorescence intensities of Sytox stained cells at different time points are shown in Additional File 1: Figure S2. Overall, these results supported that cilia motion was a crucial component for wound repair process in *Stentor*.

Second, we tested the effect of KI treatment. In untreated cells, poking the cell with a glass needle induced rapid cell contraction within ∼20 ms [20], [21], [23], [30]. We verified that KI-treated cells failed to contract upon poking, suggesting that myoneme contraction was successfully inhibited (Additional Files 10 – 11: Movies S3a – b). The survival rate of KI-treated cells was very low – about 25% for unwounded control cells and 0% for Regime 1 and 2 cells (Figure 4H), which precluded the use of our wound repair assay on KI-treated cells (all cells were washed following the KI treatment). The membranellar band of unwounded KI-treated cells remained active and beat with a mean wave propagation speed of ∼0.91 ± 0.20 mm/s and cilia beat frequency of ∼21.0 ± 5.2 Hz (N=6 cells), which was not significantly reduced compared to unwounded and untreated control cells (wave speed p=0.105; beat frequency p=0.47). In KI-treated cells wounded in Regime 2, the membranellar band beat with a mean wave propagation speed of ∼1.18 ± 0.60 mm/s and cilia beat frequency of ∼21.9 ± 4.0 Hz (N=6 cells). Compared with the smooth edges in the bands in the 2D autocorrelation of untreated control cells indicating high coordination, the bands of KI-treated cells had rough, fuzzy edges and discontinuities, indicating a slight loss of coordination (Figure 4B – C, Additional Files 12 – 13: Movies S2e – f). We observed that KI treatment intended to inhibit the myonemes could also affect the membranellar band. In about 45% of KI-treated cells, the membranellar band would be inhibited and no beating was noticeable by eye (3 biological replicates, N=18-24 cells per replicate). This inhibition could be due to the partial anesthetization of the cilia by potassium ions [31]. Iodide ions may also play a role in membranellar band inhibition, as NaI has been previously reported to inhibit the membranellar band in *Stentor*, although KI was not discussed [31]. However, the body cilia of KI-treated cells qualitatively remained as active as untreated control cells, compared to the inactive body cilia of NiCl_2_-treated cells (N=6 cells) (Additional File 14: Movie S2g). Figure 4D shows that the timescales for all three modes of response in KI-treated cells were insignificantly different from the untreated controls (contraction p=0.86; retrieval p=0.13; twisting/pulling p=0.99). In both Regime 1 and 2, KI-treated cells more often utilized contraction alone in their wound response (Regime 1, N=16/25 cells; Regime 2, N=8/27 cells) compared to untreated control cells (Regime 1, N=10/26 cells; Regime 2, N=3/31 cells), while both the cytoplasm retrieval and twisting/pulling modes occurred less often (Figure 4E). These results indicate that active myonemes are not necessary for the contraction repair mode, but may play a role in the cytoplasm retrieval and twisting/pulling modes.

## 3. Discussions

### 3.1 Timescale of wound healing

We have characterized the wound repair time in *Stentor* subject to mechanical wounds. We estimated that the wound repair rate and the maximum wound size that can be repaired in *Stentor* were both among the largest in the single cells studied thus far (Table 1). Comparison of the timescales to heal in Regimes 1 and 2 revealed that *Stentor* took longer to heal when the wounding was more severe. The timescale of each of the three mechanical modes of wound response varied from ∼20 to ∼250 seconds (Figure 4D), much shorter than ∼600 seconds for ∼90% of the cells to heal in Regime 1 (Figure 1G). We attribute this difference to the requirement of additional modes of wound repair to fully close the wound, such as vesicle trafficking and fusion with the membrane, and possibly actomyosin purse string, which may be conserved in *Stentor* due to its use in ciliate cell division [32], [33], [34]. Ongoing work is being performed to investigate these additional modes of repair, but are outside the scope of this paper.

In addition, we note that the variation in the timescale for cells to heal was relatively large even in Regime 1 (from 100 seconds where ∼50% of the cells were healed, to 600 seconds where ∼90% of the cells were healed). This variation could be due to the variation in the initial cell size (Additional File 1: Figure S4) and the wound size created by our wounding method, which relied on cell deformation and was thus sensitive to initial cell size. In addition, while the long axis of the cell was always aligned parallel to the flow and the guillotine always bisected the cell longitudinally, we could not control the orientation of the cell and the exact location where the cell was cut. This factor could lead to different types of wounds even in Regime 1, and therefore the relatively large variations in the wound healing time.

### 3.2 Mechanical modes of wound response

We have described and begun to characterize three distinct mechanical modes of wound response: contraction, cytoplasm retrieval, and twisting/pulling (Figure 5). Up to 95% of wounded cells utilized at least one of these mechanical modes, which suggests that the three mechanical modes play an important role in the wound response. While events local to the wound site have been the focus of many single-cell wound repair studies [1], the mechanical modes described here indicate that large-scale mechanical events such as cellular force generation and motility may be of greater importance than previously considered. Interestingly, no cell was observed to use all three mechanical modes. It is unknown if this is a consequence of the type and the specific location of wounds the guillotine inflicts upon the cells, or perhaps if one to two mechanical repair modes are generally sufficient for the wound repair process. In untreated Regime 2 cells, ∼90% of cells used mechanical modes of wound repair (N=28/31 cells) while only ∼70% survived (3 biological replicates, N=22-34 cells per replicate) on average. This suggests that mechanical modes of wound repair may not be sufficient for survival.

**Figure 5.**
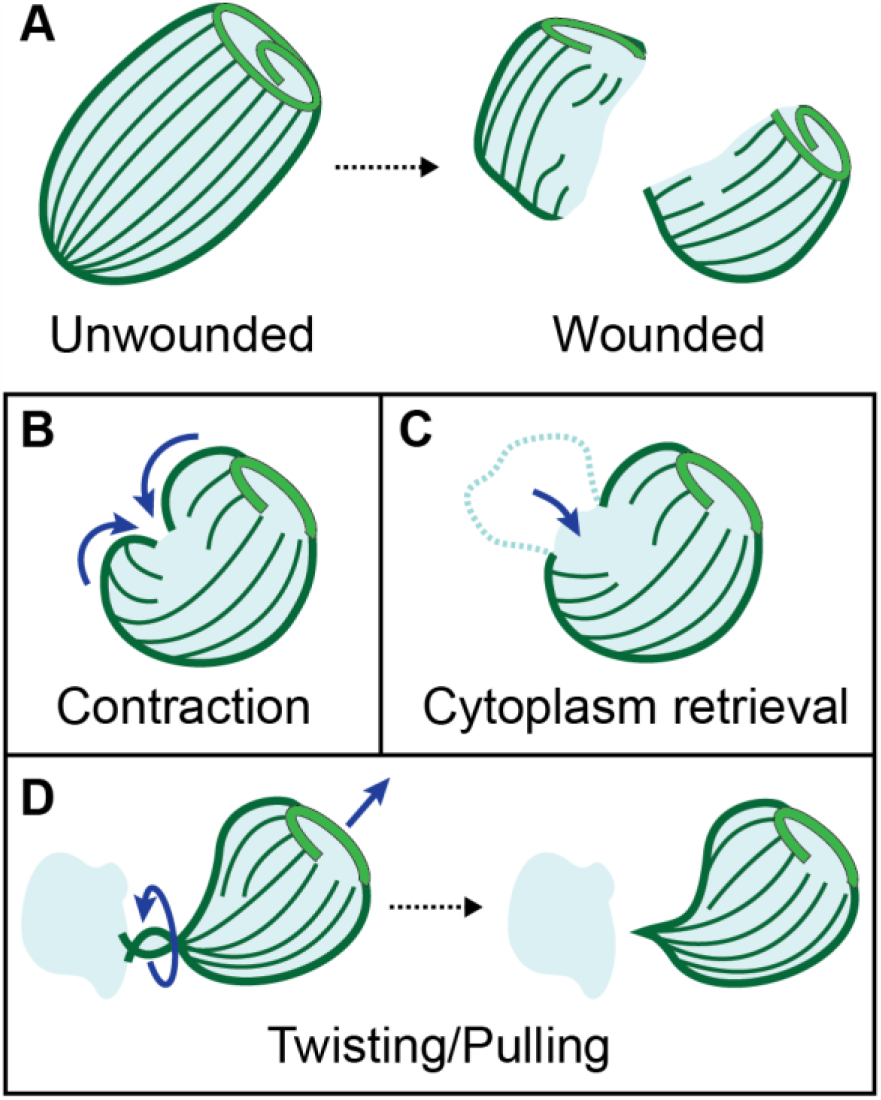
Overview of mechanical repair modes in *Stentor*. Conceptual illustration of mechanical repair modes in *Stentor* described in this study. **A)** An unwounded cell is bisected into 2 wounded cells using our microfluidic guillotine platform. After wounding, *Stentor* can potentially use a combination of 3 mechanical repair modes to aid in its wound response: **B)** contraction, in which the cell folds around the wound site, **C)** cytoplasm retrieval, in which extruded cytoplasm is pulled back into the cell body, and **D)** twisting/pulling, in which the cell uses a twisting/pulling motion to detach from extruded cytoplasm. Further work is needed to probe the signaling pathways and mechanisms of these repair modes and if *Stentor* possess any additional wound repair mechanisms.

Tartar has previously reported cells folding on themselves after being cut longitudinally [23] but no further investigation was performed. Such folding behavior was very similar to the contraction mode we had observed here. The lack of effect on the contraction by NiCl_2_ and KI treatment indicate that the process was not mediated by cilia motion or myoneme contraction. More work is necessary to investigate the mechanism underlying such contraction behavior.

In the case where the cytoplasm was extruded outside of the cell, we observed two different responses: cytoplasm retrieval and twisting/pulling. Cytoplasm retrieval has been previously reported in *Stentor* [35]. and was accompanied by extensive cytoplasmic flow that is also observed in unwounded cells. It is unknown whether this cytoplasmic flow is also driving the cytoplasm retrieval as observed here. Alternatively, it is possible that the retrieval is similar to the retraction of a bleb [36–38], where membrane and cortex reform at the boundary of the extruded cytoplasm. The contraction of the cortex then leads to the retraction of the cytoplasm inside the cell. To further probe this mechanism will require a better imaging method to verify the presence of membrane and cortex.

The twisting/pulling repair mode involved the “amputation” of the extruded cytoplasm by cilia-mediated twisting/pulling motions. Wounding did not increase the cilia beating frequency and thus the twisting/pulling mode was likely a spontaneous response, rather than a response that is activated from wounding. We note the remarkable similarity between the twisting wound response and rotokinesis, where the rotation of one half of the dividing cell severs the bridge connecting to the other cell half [33]. Rotokinesis is required for the completion of Tetrahymena cell division [33] and has also been seen in *Stentor* cell division. Without cilia motion in Tetrahymena, 60% of the mutant cells failed to complete cytokinesis [33]. Here, *Stentor* treated with NiCl_2_ had suppressed cilia motion and failed to detach from the extruded cytoplasm and heal by twisting/pulling motions. The survival of NiCl_2_-treated cells also plummeted in Regime 2 where twisting/pulling was a dominant response in healing.

The survival of NiCl_2_-treated cells in Regime 1 also decreased, but not as sharply as in Regime 2. Further, the typical wound repair time increased by an order of magnitude to ∼1000 s. This was unusual considering that the twisting/pulling repair mode was rare in Regime 1 and that the survival of the control NiCl_2_-treated cells was 100%. A possible explanation is that Ni^2+^ ions have other inhibitory effects. Ni^2+^ ions are known to inhibit axonemal dynein in *Paramecium* [29]. It is possible that cytoplasmic dynein is also affected in *Stentor*, but it is currently unclear what role cytoplasmic dynein plays in the wound repair process in *Stentor*.

KI-treated cells utilized cytoplasm retrieval and twisting/pulling less often than control cells, but the timescales of each mode were not significantly different from untreated control cells, suggesting a relatively minor role of myoneme contraction in these two modes. It is also possible that more heavily injured KI-treated cells (e.g. cells with spilled cytoplasm that would otherwise utilize cytoplasm retrieval and/or twisting/pulling) were more likely to die before a wound response could be observed, and thus be excluded from the experimental results. Although KI treatment had only a slight effect on the mechanical repair modes, the survival of wounded KI-treated cells and even the control KI-treated cells was very low, ∼0 and 25%, respectively. From these results, it is likely that KI treatment is inhibiting additional cellular processes beyond myoneme contraction, or that KI is generally cytotoxic to *Stentor*. Nevertheless, it was worthwhile to test the effects of KI treatment on the immediate wound repair response, as currently no suitable alternative is known to robustly inhibit myoneme contraction without significantly affecting cilia function, and the composition of the myonemes is not yet fully identified [22, 31].

The ability of *Stentor* to survive large open wounds for extended periods of time is an enormous biological advantage, e.g. in surviving predator attacks, and a remarkable biophysical phenomenon. Two features of *Stentor* that may contribute to this ability are of notable interest. The first is the contractile vacuole complex which periodically expels excess fluid and ions from the cell [39], a function that may be crucial in maintaining homeostasis in the presence of an open wound. The second is the massive length scale of *Stentor* cells. Because of their large size, *Stentor* may have relatively larger amounts of material reserves available to mount a wound response. Further, cell damage resulting from diffusive loss of essential biomolecules and influx of excess ions will occur on orders of magnitude slower timescales than in small cells only a few μm in size.

## 4. Conclusions

In summary, we have quantified the healing time and identified three mechanical modes of wound response in *Stentor* inflicted with mechanical wounds from a microfluidic guillotine. At least one mechanical mode was observed in almost all injured cells, which highlights the role of large-scale mechanical behaviors that may be crucial to single-cell wound repair and have not been reported in previous studies. Chemical perturbations revealed the critical role of cilia-mediated twisting/pulling motion in wound healing, and that myoneme contraction was marginally involved in the three mechanical modes observed. Future work will include further investigation of the molecular mechanisms of the wound repair process. In order to narrow the variation in cell size and wound size, future investigations could include a cell sorter to ensure the cells entering the guillotine are of uniform size.

## 5. Methods

### 5.1 *Stentor* cell culture

*Stentor coeruleus* were cultured in Pasteurized Spring Water (PSW) (132458, Carolina Biological Supplies) in 400 mL Pyrex dishes in the dark at room temperature. In a modification to standard *Stentor* culture [40], *Stentor* were fed *Chlamydomonas* every 2 days (see Additional File 1: Note S1 for details). Prior to each experiment, healthy adult cells (∼400 μm in diameter and dark green in color) were retrieved from culture by pipetting under a stereoscope into a 4 mL glass vial. A cell suspension (density ∼0.5 cells per μL) was obtained by sucking cells from the glass vial into tubing attached to a syringe filled with cell media (PSW). Cells were injected into the microfluidic guillotine for wounding experiments.

### 5.2 Microfluidic guillotine design, fabrication and wounding experiments

The master mold of the microfluidic device was fabricated in SU-8 on a silicon wafer using standard photolithography. The height of the master was measured with a profilometer to be 100 μm. A second replica mold of the master mold was made from Smoothcast 310 using a technique described previously [41]. Poly(dimethylsiloxane) (PDMS) (SYLGARD-184™, Dow Corning) was then cured from either the original or secondary mold, following standard soft lithography procedures, and bonded to a glass slide to form the final device. The device was left overnight at 65° C in an oven prior to use in order to strengthen the adhesion between PDMS and glass and to make the channel hydrophobic. Prior to use, the channels were washed with a small amount of ethanol in order to remove air bubbles from the channel. Afterwards, at least 1 mL of cell media was injected to clean the channels. Given the dimensions of our channels, the use of less than 1 mL of cell media could lead to improper cell cutting. Channels were washed from the outlet with cell media between separate experiments and discarded if cell debris or residues could not be removed from the channel.

A syringe pump was used to inject the cell suspension into the microfluidic guillotine (inlet **b**, Figure 1A – B) at constant flow rates of 8 mL/hr for Regime 1 and 36 mL/hr for Regime 2 respectively. These flow rates corresponded to an average velocity of 1.4 cm/s and 6.3 cm/s respectively at each guillotine. Cell media was injected *via* inlet **a (**Figure 1A – B) at various flow rates to control the wound repair time (see Section 5.3).

### 5.3 Controlling wound repair time prior to fixation and staining

We controlled the duration between the time when cells were wounded and when they were fixed (t_post-wound_) using two methods.

- 1) t_post-wound_ < 30 seconds. We varied the length of the outlet tubing (BB31695-PE/4, Scientific Commodities Inc.) from 4 cm to 25 cm with a constant cross-section area of 0.454 mm^2^.
  ○ The cell suspension was injected into the device via inlet **b** at the flow rates listed in Section 2.2. In Regime 1, cell media was also simultaneously injected *via* inlet **a** at 10 mL/hr to increase the velocity of cells exiting the outlet **c** to 18 mL/hr total, so to decrease the lag time before assaying the cells. Increasing the flow rate at the outlet also prevented cells from sticking to the tubing, increasing the accuracy of t_post-wound_.
  ○ Additional File 1: Note S2 details how t_post-wound_ was calculated.
- 2) 1 minute < t_post-wound_ < 120 minutes. We varied the incubation time of the cells inside the tubing. The outlet tubing length was standardized (25 cm) and was prefilled with cell media and went into an empty 2 mL-Eppendorf tube. A constant volume of cell suspension (110 μL for Regime 1, and 70 μL for Regime 2) was injected *via* inlet **b** into the microfluidic guillotine device at the flow rates listed in Section 2.1. Note that inlet **a** was not used at this time, but was connected to a syringe to prevent backflow into inlet **a**.
  ○ After the fixed volume of cell suspension was injected, a timer was started immediately. The majority of wounded cells were inside the outlet tubing, and cells that remained in the device were not fixed or stained.
  ○ When the timer went off, corresponding to the desired incubation time prior to fixation and staining, we pumped 250 μL of cells into 1 mL of fixing solution from the flushing inlet (inlet **a**) at 18 mL/hr (for both Regime 1 and 2 experiments). By flushing with inlet **a**, which went directly to the outlet, we omitted any cells which were not already in the tubing after wounding. Note that inlet **b** was not used at this time, but had a syringe hooked up to it that prevented backflow into inlet **b**.
  ○ Additional File 1: Note S2 details how t_post-wound_ was calculated.

### 5.4 Fixation protocol and wound repair assay

Extreme care was taken in all steps to minimize cell wounding during handling and fixation. We used a fixing solution of 1% formaldehyde (no methanol) (43368, Alfa Aesar) and 0.025% Triton X-100 (X100-100ML, Sigma Life Sciences) in PSW at room temperature. To fix the cells, 250 μL of wounded cells was ejected from the guillotine device and the outlet tubing using a syringe pump (see details in Section 5.3) into 1 mL of the fixing solution in a 2 mL round-bottomed tube (111568, Globe Scientific) and incubated for 10 minutes at room temperature. The round bottom aided in the collection of cells while avoiding significant clumping. The tubing was submerged in the fixation solution and did not contact the bottom of the 2 mL tube to avoid additional cell wounding.

To stain the wounded cells, we used Sytox Green (S7020, Invitrogen) at a concentration of 2.5 μM in PSW at room temperature. After fixation, we used a 200 μL pipette tip with the end cut off at the 20 μL line to transfer 50 μL of the cells from the bottom of the fixation tube to a 500 μL solution of Sytox in a 4 mL glass vial with a flat bottom (C4015-21, ThermoScientific). A flat bottom vial, rather than a rounded or conical bottom vial, was used to ensure all cells were exposed evenly to Sytox as cells did not pellet in the flat bottom vial. After 30 minutes of incubation in Sytox, we washed 50 μL of the stained cells in 500 μL of PSW in a second 4 mL glass vial, and then transferred 50 μL of washed cells onto a No. 1 glass slide using a 200 μL pipette tip with the end cut off at the 20 μL line. If we could not capture all cells in the 50 μL volume, additional 50 μL volumes were put onto separate glass slides to avoid wounding the cells already on the slide and to avoid increasing the background signal due to additional media volume on the slide. To minimize cell adhesion and damage due to shear, tubing and pipette tips used to handle wounded cells were treated with 3% Pluronic F-68 (J6608736, Alfa Aesar) in deionized (DI) water for 3 hours and then washed.

Cells were then imaged at 15x magnification on an EMCCD camera (Andor iXon 897, Oxford Instruments) mounted on an epifluorescence microscope with 0.05 s exposure time, a mercury lamp set to ND 1, and a FITC excitation / emission filter set. To quantify the fluorescence of cells stained with Sytox, we manually traced the cells in ImageJ and measured the average pixel intensity of each cell. See Additional File 1: Figure S5, Note S3 for the complete reagent list and for the details on the optimization of the assay.

### 5.5 Immunostaining of acetylated tubulin to visualize KM fibers

To visualize KM fibers in wounded and unwounded cells, we performed immunofluorescence for acetylated tubulin. To begin, we fixed 250 μL of cells in 1000 μL of ice-cold methanol in a 2 mL plastic tube (16466-060, VWR). For wounded cells, the time post-wounding was controlled following the methods in Section 5.3. Cells were incubated in methanol at -20 °C for 40 minutes. We removed the supernatant and then added 500 μL of 1:1 methanol:PBS and incubated for 10 minutes at room temperature. We removed the supernatant and then added 500 μL of PBS and incubated for 20 minutes at room temperature. We removed the supernatant and then and added blocking buffer consisting of 500 μL of 2% BSA + 0.1% Triton X-100 in PBS and incubated for 2 hours at room temperature or overnight at 4 °C. We removed the supernatant and then added 500 μL of primary antibody (T7451-200UL, Sigma Life Sciences) diluted 1:1000 in blocking buffer and incubated for 2 hours at room temperature or overnight at 4 °C. We removed the supernatant and then washed 3 times with 500 μL of PBS, allowing cells to settle at the bottom of the tube between each wash. We removed the supernatant and then added 500 μL of 488 nm excitation fluorescent secondary antibody (SAB4600388-125UL, Fluka Analytics) diluted 1:1000 in blocking buffer and incubated for 2 hours at room temperature or overnight at 4 °C. We removed the supernatant and then washed 3 times with 500 μL of PBS, allowing cells to settle at the bottom of the tube between each wash. We then immediately imaged the cells by pipetting 50 μL onto a No. 1 glass slide.

We used two imaging setups for the KM fibers. We obtained confocal images using an inverted laser scanning confocal microscope (Zeiss, LSM 780). Cells were imaged using a 20X (NA = 0.8) objective at an excitation wavelength of 488 nm, and a broad emission filter matching the spectra of Alexa Fluor 488. We obtained additional epifluorescence images using an EMCCD camera (Andor iXon 897, Oxford Instruments) mounted on an inverted microscope with a 10x or 20x objective.

### 5.6 Imaging of cell motion after wounding

We observed cell repair behavior using a high-speed camera (Phantom v7.3 or Phantom v341) operating between 20 - 100 fps mounted on an inverted brightfield microscope using a 4x, 5x, or 10x objective. Cell repair behaviors were observed immediately after cutting, either inside the device in the large well before the outlet, or on top of the PDMS device using No. 1.5 coverslips to create a well with a defined height.

To measure the time for each mechanical mode to complete, we determined the first and last frame the mode could be seen manually. The contraction mode was considered complete when the wound was smaller than ∼20 μm, at which point it was difficult to see consistently with our imaging setup. The twisting/pulling modes were considered complete when the cell completely detached itself from the extruded cytoplasm. The cytoplasm retrieval mode was considered complete when the cytoplasm did not form an apparent protrusion out of the cell.

To measure the timescale and frequency of occurrence of different modes of wound response, we only considered cells and behaviors which met the following criteria: 1) The location of wounding was visible with our imaging setup. For example, the wound location would not be visible if it were obscured by the shadow from the device sidewalls or the outlet tubing, or if the cell left the field of view. If we lost visibility of the wound location before the wound response was complete, we would count behaviors that were observed prior to the loss of visibility towards the frequency of occurrence, but would not quantify the timescale of their response. 2) The cell did not swim back into the narrow channels near the guillotine blade during its wound response. Re-entry into these narrow channels would deform and compress the cells, possibly leading to additional injury. If a cell entered the narrow channels before its wound response was complete, we would count behaviors that were observed prior to the contact towards the frequency of occurrence, but would not quantify the timescale of their response. 3) Unwounded cells (due to stopping of the pump for observation) were not included. 4) The cell did not die during observation (which typically occurred in the first 5 – 10 minutes after wounding). For this experiment, we considered cells dead if they ruptured and lost complete membrane integrity. This type of cell death was only observed for a few cases with KI treatment.

The frequency of each behavior was then calculated by considering cells which met all criteria for that behavior. Some cells used multiple modes, and thus the frequency of occurrence for the three modes can add up to greater than 100%. For cases where the mechanical healing modes were less common, the sum of the frequencies of occurrence can be less than 100%, indicating that not all wounded cells utilized a mechanical healing mode.

### 5.7 Survival measurements

Cell survival after cutting was quantified using Eq. 1:

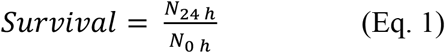

where *N*_0 *h*_ is the number of cell fragments counted at the channel outlet at t = 0 hours, immediately after the cut, and *N*_24 *h*_ is the number of live cells 24 hours after the cut. *N*_0 *h*_ was counted manually from videos of the cutting process. The videos were obtained using the high-speed camera at 100 fps (Regime 1) and at 450 fps (Regime 2) respectively. Immediately after cutting, cell fragments were included in *N*_*0 h*_ if they were larger than 1/4 of a typical bisected cell cut in Regime 1 and appeared to have a part of an intact cell membrane. *N*_24 *h*_ was counted manually by examining the cells stored overnight in a 2” Petri dish with ∼5 mL of cell media. At 24 hours after cutting, cells were considered alive and included in *N*_*24 h*_ if they had beating cilia, were swimming, or were attached to the surface in a trumpet-like shape[24].

### 5.8 Perturbations

#### 5.8.1 Nickel chloride (NiCl_2_) treatment

Cilia beating was reversibly inhibited by NiCl_2_ treatment. We prepared a 100 mM stock solution of NiCl_2(aq)_ (N6136, Sigma) in DI water and gently heated it at 50° C until fully dissolved.

Inhibiting cilia beating caused increased adhesion of the cells to the tubing and the microfluidic device. To minimize cell adhesion and damage due to shear, all tubing and microfluidic devices used in the NiCl_2_ experiments were incubated with 3% Pluronic F68 in DI water for 3 hours and then washed.

To prepare NiCl_2_ treated cells, we loaded approximately 15 cells in a 25 μM NiCl_2_ solution in PSW inside a 20 cm length of tubing using a syringe. The tubing was then disconnected from the syringe (to prevent cells from swimming into the syringe barrel before the cilia were fully inhibited) and the cells were incubated for 5 hours. Then, the tubing was re-connected to a syringe and prepared device both filled with cell media, taking care to eliminate any air bubbles. From this point onwards, device operation, cell fixing, and Sytox staining proceeded as before. We found this treatment condition was the most effective in inhibiting cilia beating without compromising cell viability. Cilia motion was recovered within 24 hours post-wash.

#### 5.8.2 Potassium iodide (KI) treatment

We incubated cells in a solution of 1% KI (60400, Sigma-Aldrich) in PSW for 5 minutes, followed by washing in PSW. Cells were used within 30 minutes after washing. This method has been shown previously to inhibit *Stentor* contraction [23]. To minimize cell adhesion and damage due to shear, all tubing and microfluidic devices used in the KI experiments were incubated with 3% Pluronic F68 in DI water for 3 hours and then washed.

### 5.9 Measurement of cilia activity

High-speed images of the cells were taken at 5000 fps with a 20x objective using the Phantom v341 camera and analyzed as described previously[42]. Briefly, a 1D coordinate system along the membranellar band was defined using ImageJ and used to construct a region of interest. 2D kymographs of image intensity were computed and autocorrelated to determine the spatiotemporal coordination of the beating cilia following the method published by Wan et al.[42]

### 5.10 Experimental design and statistical analysis

Cells used for experiments were always used the second day after feeding to ensure consistency (Additional File 1: Notes S1, S3). At least 3 biological replicates were performed for each experimental group in the wound repair assay experiments. For wound repair mechanism observation experiments, each independent experiment yielded 1-3 cells for observation, and a total of at least 25 cells were observed for each experimental group. To assess changes in wound repair mechanism timescales under drug treatment, we performed statistical analysis using unpaired two-sample t*-*tests with Holm-Sidak correction for multiple comparisons. Values of p< 0.05 were considered statistically significant. The sample size N in each experiment group is provided in the figure captions. The same batch of reagents were used throughout all experiments to minimize batch to batch variation.

## Supporting information

Additional_File_1_FiguresNotes

Additional_File_2_Movie_S1a_Contraction

Additional_File_3_Movie_S1b_CytoplasmRetrieval

Additional_File_4_Movie_S1c_Twisting

Additional_File_5_Movie_S1d_Pulling

Additional_File_6_Movie_S2a_cilia_PSW_unwounded

Additional_File_7_Movie_S2b_cilia_PSW_wounded

Additional_File_8_Movie_S2c_cilia_NiCl2_unwounded

Additional_File_9_Movie_S2d_cilia_NiCl2_wounded

Additional_File_10_Movie_S3a_control_contraction

Additional_File_11_Movie_S3b_KI_contraction

Additional_File_12_Movie_S2e_cilia_KI_unwounded

Additional_File_13_Movie_S2f_cilia_KI_wounded

Additional_File_14_Movie_S2g_BodyCilia_unwounded

## 6. Declarations

### Ethics approval and consent to participate

Not applicable

### Consent for publication

Not applicable

### Availability of data and materials

The datasets used and/or analyzed during the current study are available from the corresponding author on reasonable request.

## Authors’ contributions

KSZ, LRB, SKYT designed the experiments; KSZ, LRB, WH performed experiments and data analysis. KSZ, LRB, WFM, and SKYT wrote the manuscript.

### Acknowledgements

We thank the instructors and students of the 2019 Center for Cellular Construction Summer Course for their helpful discussions.

## Additional Files

**Additional_File_1_FiguresNotes.docx**

Contains the following supplementary figures and notes:

Figure S1. Additional images of cells post wounding

Figure S2. Measured mean fluorescence of individual cells using the wound repair assay

Figure S3. Details of cell behavior quantification

Figure S4. Cell size variation

Figure S5. Optimization of Sytox Green staining

Note S1. *Stentor* feeding protocol, optimized for wound repair assay.

Note S2. Calculation of wound repair time.

Note S3. Wound repair assay optimization

Table S1. Comparison of different dyes for staining wounded *Stentor* cells.

**[[Movie set S1: Mechanical wound responses]]**

All movies were taken at 20 fps with a 10x objective using the Phantom v7.3 camera.

**Additional_File_2_Movie_S1a_Contraction**.**mp4**

Observation of the contraction wound response in Regime 1.

**Additional_File_3_Movie_S1b_CytoplasmRetrieval**.**mp4**

Observation of the cytoplasm retrieval wound response in Regime 2.

**Additional_File_4_Movie_S1c_Twisting**.**avi**

Observation of the twisting wound response in Regime 2.

**Additional_File_5_Movie_S1d_Pulling**.**mp4**

Two observations of the pulling wound repair in Regime 2.

**[[Movie set S2. Cilia velocity in different perturbations]]**

Movies S2a – S2f in Movie set S2 are representative of the cilia motion at the membranellar band for their respective experimental condition. Movie S2g is representative of body cilia motion. All movies were taken at 5000 fps with a 20x objective using the Phantom v341 camera except for Movie S2g, which was taken at 1000 fps with a 40x air objective.

**Additional_File_6_Movie_S2a_cilia_PSW_unwounded**.**mp4**

**Additional_File_7_Movie_S2b_cilia_PSW_wounded**.**mp4**

**Additional_File_8_Movie_S2c_cilia_NiCl2_unwounded**.**mp4**

**Additional_File_9_Movie_S2d_cilia_NiCl2_wounded**.**mp4**

**[[Movie set S3. KI inhibition of myoneme contraction]]**

KI inhibition of myoneme contraction demonstrated by poking cells with a glass needle. Movies were taken at 450fps with a 5x objective using the Phantom v341 camera.

**Additional_File_10_Movie_S3a_control_contraction**.**avi Additional_File_11_Movie_S3b_KI_contraction**.**avi**

**[[Movie set S2. Cilia velocity in different perturbations]] (Continued)**

**Additional_File_12_Movie_S2e_cilia_KI_unwounded**.**mp4**

**Additional_File_13_Movie_S2f_cilia_KI_wounded**.**mp4**

**Additional_File_14_Movie_S2g_BodyCilia_unwounded**.**mp4**

