## Additional_File_1_FiguresNotes for "Microfluidic guillotine reveals multiple timescales and mechanical modes of wound response in *Stentor coeruleus*"

**Additional File 1** – Supplementary figures and notes

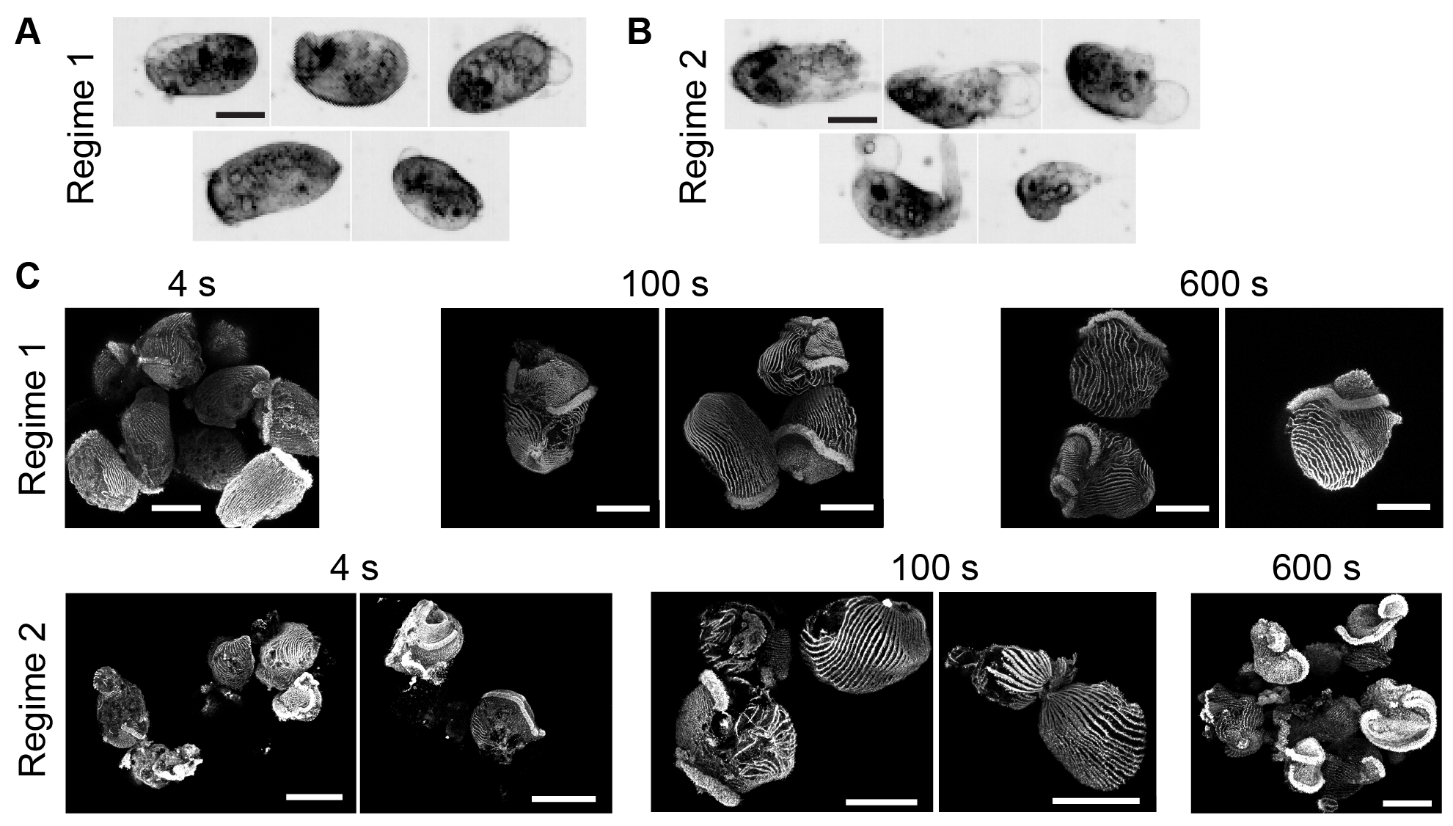

**Figure S1. Additional images of cells post wounding**

**A)** Brightfield images of cells cut in Regime 1. All images were taken within 1 min of wounding. **B)** Brightfield images of cells cut in Regime 2. All images were taken within 1 min of wounding. **C)** Additional immunostaining images of KM fibers in cells. Cells were wounded in either Regime 1 or Regime 2 and fixed at 4 s, 100 s, or 10 min after wounding. These images were acquired on the inverted confocal microscope. All scale bars 100 μm.

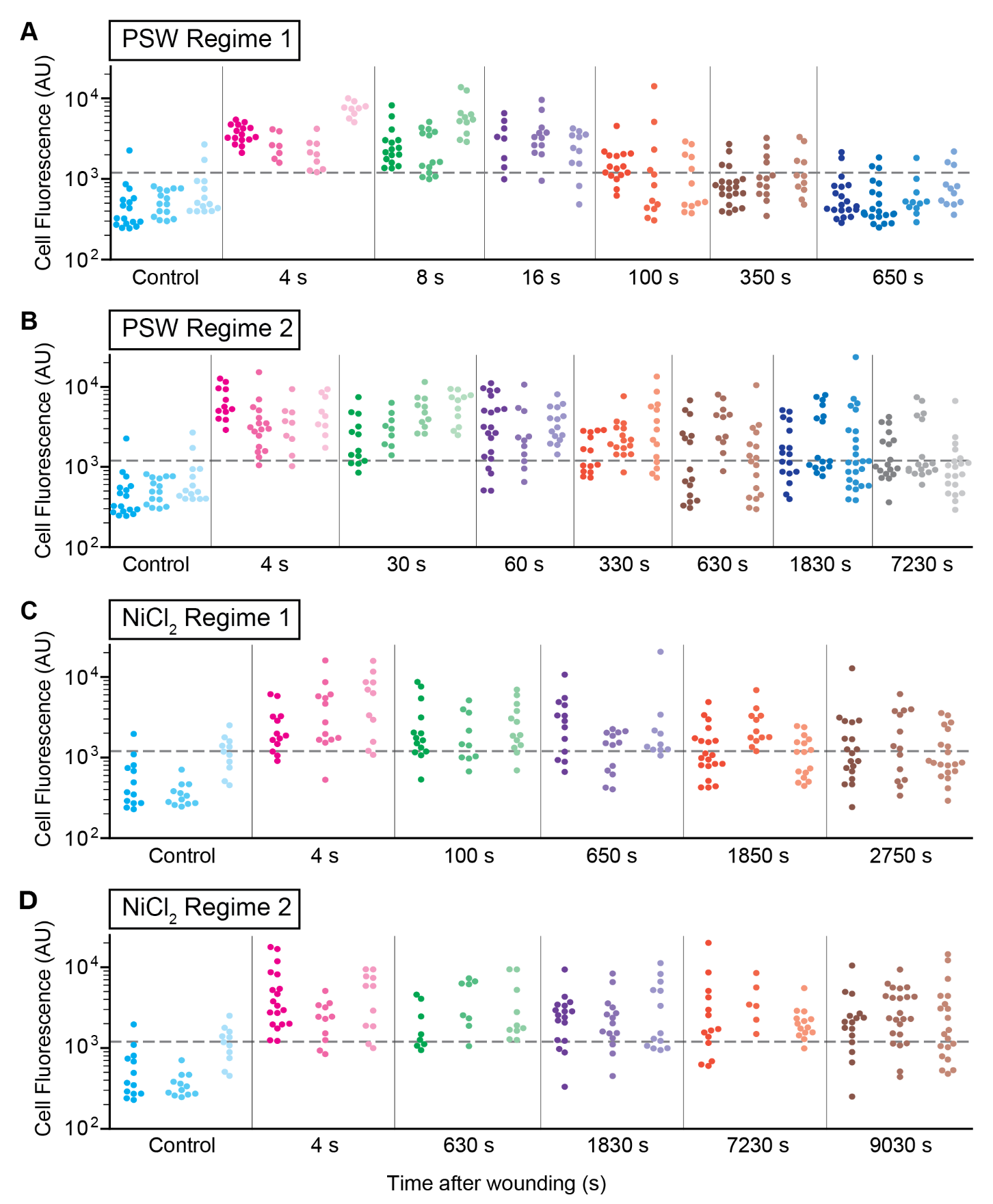

**Figure S2. Measured mean fluorescence of individual cells using the wound repair assay**

Measured mean fluorescence intensity values for individual cells (dots) across all experimental conditions and biological replicates, using the wound repair assay. At least 3 biological replicates were performed for each experimental condition, represented as individual swarmplots. Dotted line indicates I­_threshold_ = 1200 AU used to distinguish unwounded or healed cells from wounded cells. **A)** PSW (untreated) cells wounded in Regime 1 (N=7-22 cells per replicate). **B)** PSW (untreated) cells wounded in Regime 2 (N=9-25 cells per replicate). **C)** NiCl_2_-treated cells wounded in Regime 1 (N=11-20 cells per replicate). **D)** NiCl_2_-treated cells wounded in Regime 2 (N=11-23 cells per replicate).

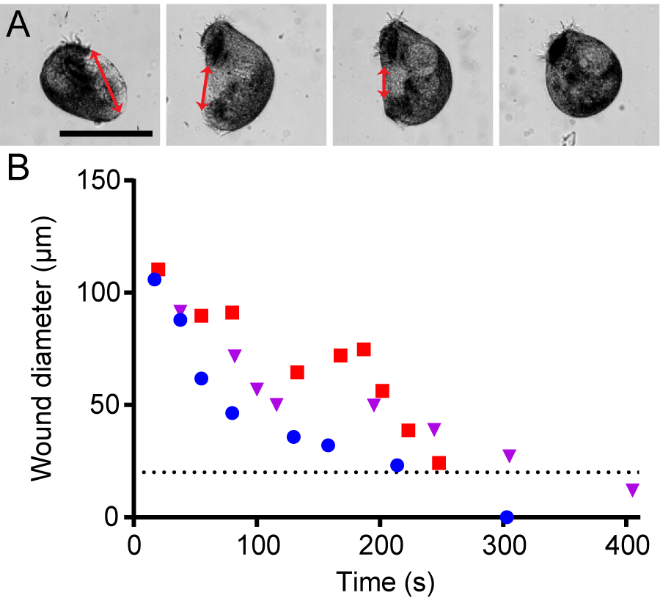

**Figure S3. Details of cell behavior quantification**

**A)** The wound diameter was estimated by drawing a line through the wound as shown. **B)** Wound diameter over time for three example cells cut in Regime 1 that used the contraction mode. The cells were said to be healed when their wound was approximately equal to 20 μm, shown by the horizontal line. At 20 μm, the wound was only around 10 pixels in diameter and was difficult to see consistently in all cells.

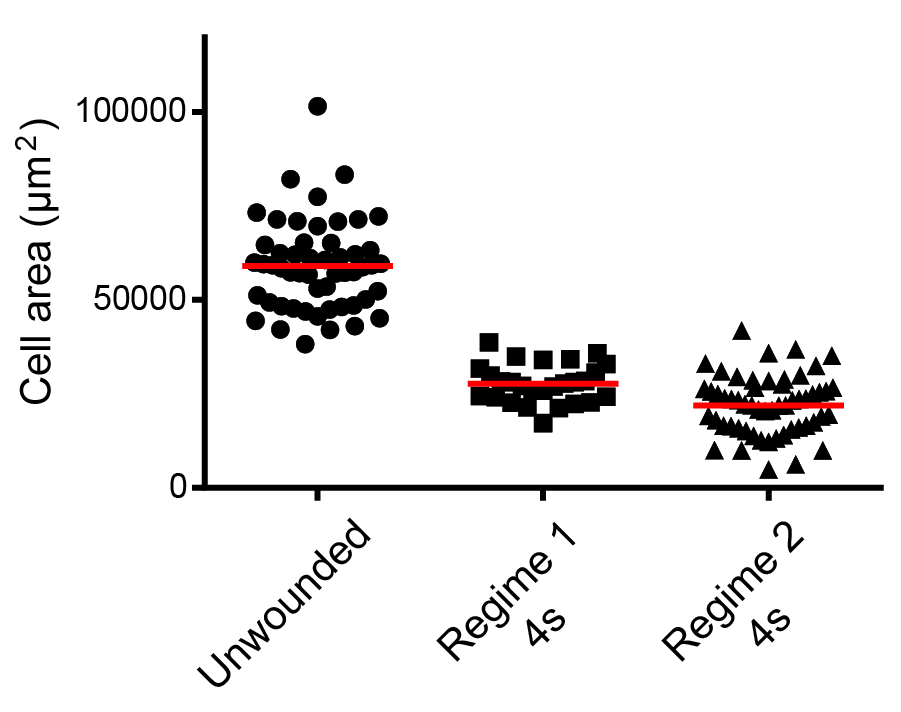

**Figure S4. Cell size variation**

Cell area for unwounded cells and cells cut in Regimes 1 and 2s, 4 s after wounding. Red line represents the mean of the data. Cells were fixed following the wound repair assay protocol prior to imaging with an EMCCD camera at 10x magnification. N=27–56.

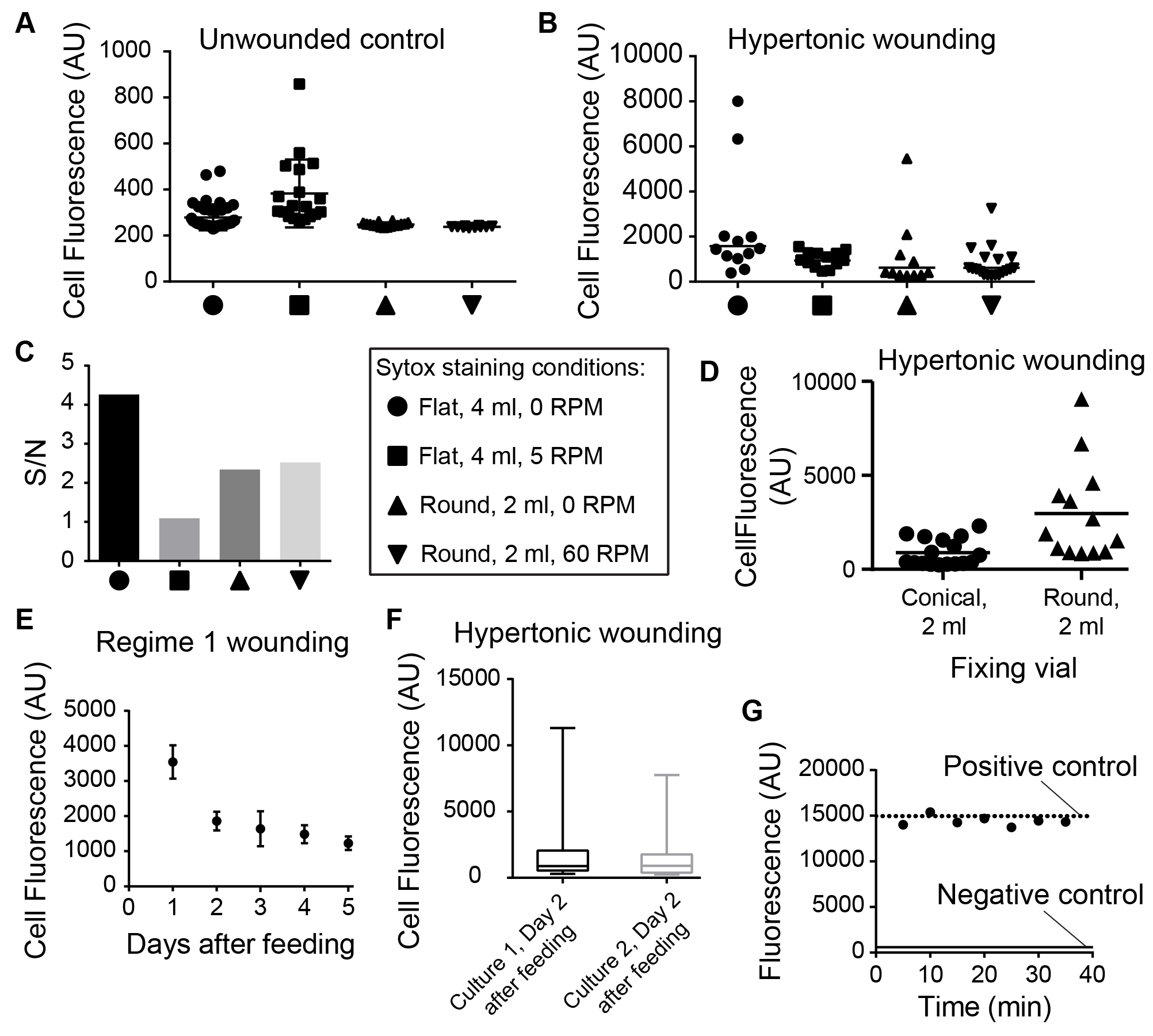

**Figure S5. Optimization of Sytox Green staining**

**A)** Sytox Green fluorescence for unwounded cells incubated in Sytox in different tubes. Flat and round refer to the shape of the bottom of the tube. We also tested the use of an orbital shaker to increase mixing. **B)** Sytox Green fluorescence for cells osmotically wounded by fixing in a hypertonic solution of 4x HBSS in the same conditions as **A)**. **C)** Signal-to-noise ratio (S/N) of the fluorescence intensity for wounded cells compared to unwounded cells, calculated by dividing the mean fluorescence in **B)** by that in **A)**. The S/N was the highest for the flat 4 mL tube without shaking. **D)** Sytox Green fluorescence for cells fixing cells in a 2 mL tube with a conical bottom and round bottom. Cells were wounded by fixing in a hypertonic solution of 4x HBSS. **E)** Sytox Green fluorescence for cells wounded at 8 mL/hr, 4 s after wounding, as a function of days after feeding. Cells in this experiment were fed with *Chlamydomonas* on a 5 day feeding schedule. **F)** Sytox Green fluorescence for cells wounded in a hypertonic solution of 4x HBSS for two different cultures on day 2 after feeding. Cells in this experiment were fed with *Chlamydomonas* on a 2 day feeding schedule. **G)** Sytox fluorescence over time when incubated with fixing solution. The experimental condition contained 2 μL of plasmid DNA (1 mg/mL stock) in 1000 μL Sytox solution and 100 μL fixing solution. This concentration of fixing solution was slightly higher than we used in our Sytox wound repair assay. The positive control was 1000 μL of Sytox solution with 2 μL of plasmid DNA (1 mg/mL stock) and 100 μL of DI water. The negative control was 1000 μL of Sytox with 100 μL of fixing solution.

**Note S1. *Stentor* feeding protocol, optimized for wound repair assay.**

We followed a standard protocol for *Stentor* cell culture [40, main text], except for the standard 5-day feeding schedule for established cell cultures. Instead, we switched to a 2-day feeding schedule, feeding 40% of the amount normally fed to the cells on a 5-day schedule. We adjusted the amount of PSW to maintain *Stentor* cell density around 20 cells/mL and adjusted the amount of algae fed depending on the approximate number of cells in the culture. For *Stentor* at 20 cells/mL, we fed 1 mL of concentrated algae per 100 mL of culture, or 1 mL algae per approximately 2000 cells. To prepare 1 mL of concentrated algae, we spun down 2 mL of *Chlyamydomonas* (after they appeared healthy and very green) cultured under light in TAP media using a centrifuge at 2x G, removed the TAP, washed with 2 mL of PSW, repeated the centrifugation step, and washed with 1 mL of PSW. The 2-day feeding schedule was crucial for obtaining consistent results using the wound repair assay (see Additional File 1: Note S3.5).

**Note S2. Calculation of wound repair time.**

***For t_post-wound_ < 30 seconds:***

We calculated t_post-wound_ by calculating the flow velocity from the flowrate at the outlet (Regime 1: cell inlet flowrate + media inlet flowrate = 18 mL/hr. Regime 2: cell inlet flowrate = 36 mL/hr; media inlet flowrate = 0 mL/hr and was stopped with a glass plug) and the inner diameter of the cylindrical tubing (= 0.76 mm), followed by dividing the length of the tubing (= 4 to 25 cm) by the flow velocity.

***For 1 min < t_post-wound_ < 150 min:***

For these timepoints, we incubated the cells in the outlet tubing for a fixed time after the cells had been cut (see Section 5.3, main text), after which we started flow from inlet **a**. If a cell is cut immediately after starting flow from inlet **b**, it will be towards the end of the tubing and may immediately enter the fixative after starting flow from inlet **a**.

For Regime 1:

t_post-wound,1_ = (time to pump 110 uL at 8 mL/hr from inlet **b**) + (incubation time) = 49.5 s + (incubation time)

For Regime 2:

t_post-wound,1_ = (time to pump 70 uL at 36 mL/hr from inlet **b**) + (incubation time) = 7 s + (incubation time)

If a cell is cut at the very end of flow from inlet **a**, then it will be towards the beginning of the tubing and it may enter the fixative at the very end of flow from inlet **a**:

t_post-wound,2_ = (time to pump 250 uL at 18 mL/hr from inlet **a**) + (incubation time) = 50 s + (incubation time)

We averaged the two values to get the final healing time t_post-wound_ = (t_post-wound,1_ + t_post-wound,2_ )/2

**Note S3. Wound repair assay optimization**

Performance of the wound repair assay was difficult and requires practice and concentration. Every step should be strictly adhered to, including the cell feeding schedule and containers used during fixation and staining. Failure to do so will likely result in further optimization being required on the part of the experimenter. To determine the success of the protocol, we recommend starting with unwounded cells, cells immediately after wounding, and cells that have had at least 10 minutes to heal (for Regime 1). For these conditions, one should expect to have very low fluorescence, very high fluorescence, and very low fluorescence, respectively.

***S3.1 Fixation and wound repair assay reagent and supply list***

- 2 mL round-bottom tube for fixation (111568, Globe Scientific)
- 4 mL glass vial for Sytox staining (C4015-21, ThermoScientific)
- No. 1 glass slides for cell imaging (1418-10, Globe Scientific)
- Formaldehyde, 16% (43368, Alfa Aesar)
- Triton, 100% (X100-100ML, Sigma Life Sciences)
- Sytox Green (S7020, Invitrogen)
- 10X HBSS (14185-052, Gibco)
- Pluronic F-68 (J6608736, Alfa Aesar)
- Pasteurized Spring Water (132458, Carolina Biological Supplies)

***S3.2 Tubing connecting the guillotine and the fixing solution.***

18 mL/hr was chosen to eject the wounded cells from the tubing into the fixing solution to minimize cell sticking to the tubing, which often occurred at lower flow rates, and to minimize cell wounding at higher flowrates. We did not use LDPE tubing longer than 25 cm to transfer wounded cells, as our preliminary data suggested that longer tubing increased the Sytox fluorescence of some wounded cells. We also did not allow longer tubing to twist or curl, which we suspected could increase cell wounding due to the flow changing direction in each twist. We did not reuse tubing between days to avoid contamination or cells sticking to fouled tubing. We submerged the tubing in the fixation solution immediately prior to starting the experiment and did not allow tubing to touch the bottom of the tube to avoid additional cell wounding.

***S3.3 Fixation protocol optimization.***

In order to determine whether a fixed cell had previously healed its wounds, a fixation protocol had to be developed that does not create additional wounds to the cell. This immediately ruled out methanol and other alcohol-based fixation methods, which permeabilize the cell membrane and displace water inside the cell. Formaldehyde is a better choice for cell fixation without permeabilization, as it does not inherently permeabilize the cell. However, when testing formaldehyde fixation we discovered that unwounded cells often became wounded during the fixation process due to the cell membrane bursting open and that cells adhered to the side walls of the fixing vial, which increased the risk of damage to the cells during collection. Using a lower formaldehyde concentration and increasing the fixing time resolved these two issues without over-cross-linking wounded cells. We also observed that adding small amounts of Triton X-100 surfactant to the fixing media and treating pipette tips with Pluronic F-68 reduced cell adhesion without increasing Sytox fluorescence in unwounded cells. The final fixation media contained 1% formaldehyde and 0.025% Triton X-100 in PSW. In our final fixation protocol, we added 250 uL of cells in PSW to 1000 uL of fixation media and incubated for 10 minutes at room temperature. Adding a smaller volume of cells to a large volume of fixation media minimized the difference in osmotic pressure felt by the cells which entered the media at different times.

***S3.4 Vial size and shaking conditions.***

We wanted to maximize the signal to noise (S/N) of the wound repair assay in order to detect small wounds and increase our confidence in our measured wound repair time. In order to maximize the S/N, we used vials which would increase exposure of cells to Sytox and also tested the use of an orbital shaker to increase mixing (Additional File 1: Figure S5A – C). To measure S/N, we osmotically wounded cells at the fixation step by fixing in a hypertonic solution of 4X HBSS (Additional File 1: Figure S5B). We then compared the fluorescence to unwounded cells fixed in normal fixing media (Additional File 1: Figure S5A). We found that incubating cells in Sytox in a 4 mL glass vial with a large flat bottom helped to minimize cell clumping and increase exposure of cells to Sytox, maximizing S/N (Additional File 1: Figure S5C). We also tested light shaking and mixing of the vials, which could have increased exposure of cells to Sytox, but we found it increased the fluorescence of unwounded cells and thus lowered S/N. When fixing in conical bottom tubes, we observed cells often clumped together at the bottom of the fixing tube and would remain clumped during Sytox incubation. We found that this led to many cells having a decreased Sytox fluorescence, likely because they were not fully exposed to Sytox. We found that fixing cells in a 2 mL tube with a round bottom (RB), rather than a conical bottom, helped to increase Sytox fluorescence for wounded cells but not for unwounded cells (Additional File 1: Figure S5D). We did not wound more than 25 cells at one time to prevent cell clumping at the bottom of the tube.

***S3.5 Stentor feeding schedule.***

Since Sytox is a nucleic acid stain, its fluorescence is determined by the concentration of nucleic acids such as RNA and DNA inside the cell. We found that when we fed the cells on a 5-day schedule, as was the standard for *Stentor* cell culture [40, main text], the fluorescence of wounded cells was much higher on day 1 after feeding and then decreased day to day through day 5 (Additional File 1: Figure S5E).

We wanted to maximize dynamic range of our assay, and thus have as high fluorescence as possible, so we switched to a 2-day feeding schedule to keep the cells consistently well-fed (See Additional File 1: Note S1). We only used cells on the second day after feeding to minimize any unexpected differences between cells and Sytox fluorescence caused by cell division, which typically occurred the day after feeding.

When we performed these changes to the feeding protocol, we found that the fluorescence of wounded cells was high and consistent between cultures (P = 0.2, NS), shown in Additional File 1: Figure S5F. For this experiment we consistently wounded cells by fixing in a hypertonic solution of 4X HBSS.

***S3.6 Checking the fidelity of Sytox.***

In the data sheet for Sytox it does not recommend using it with fixed cells as Sytox could degrade. We had already found success using Sytox with fixed *Stentor*, but we tested whether Sytox fluorescence was affected over the time course of our incubation when incubating with fixation solution at the concentration we used.

To determine if fixative decreased Sytox fluorescence, we put 2 μL of plasmid DNA (1 mg/mL stock) from Andrew Fire’s lab at Stanford into 1000 μL Sytox solution and 100 μL fixing solution. This concentration of fixing solution was slightly higher than the one we used in our Sytox wound repair assay. Our positive control was 1000 μL Sytox with 2 μL of plasmid DNA (1 mg/mL stock) and 100 μL of DI water. Our negative control was 1000 μL of Sytox with 100 μL of fixing solution. Fluorescence was measured using our Andor camera and a 96 well plate with 100 uL of solution per well. Additional File 1: Figure S5G shows that we observed no change in fluorescence of the experimental condition over 35 minutes of observation, where it remained approximately equal in fluorescence to the positive control. Therefore, we conclude that Sytox fluorescence is not affected by the fixing solution over the time course of our experiment.

***S3.7 Staining for wound repair.***

The ideal wound repair dye has several important characteristics: 1. It does not penetrate the membrane of unwounded cells and does not fluoresce. 2. It penetrates the membrane of wounded cells and is highly fluorescent. 3. The fluorescence is proportional to the size of the wound. 4. It can be used in fixed cells, since live *Stentor* are highly motile and very difficult to image. 5. It has consistent fluorescence between different cells on different days. To determine the dye for our wound repair assay, we tested several different dyes to find the right dye for our wound repair assay. These dyes are listed in Additional File 1: Table S1. In preliminary experiments on fixed cells we found that Sytox Green gave excellent fluorescence signal in wounded cells and very low signal in unwounded cells. Additionally, background signal in Sytox Green is essentially non-existent, meaning that it could be used without a final washing step.

**Table S1. Comparison of different dyes for staining wounded *Stentor* cells.**

| **Dye** | **Mode of action** | **Ex / Em** | **Comments** |
| --- | --- | --- | --- |
| Sytox Green | Membrane impermeable. Binds to nucleotides in the cell. | Blue / green | Very bright in wounded cells only. Can be used in fixed cells. Does not require a washing step. Negative effects were observed on unfixed cells at concentrations high enough for signal. |
| Propidium Iodide | Membrane impermeable. Binds to nucleotides in the cell. | Green / red | Improper wavelength due to *Stentor* autofluorescence. |
| Trypan Blue | Membrane impermeable. | N.A. | Cannot easily see difference between unwounded cell and background. Higher concentrations kill *Stentor*. |
| Live/Dead Fixable Green | Membrane impermeable. Reacts to amines within the cell. | Blue / green | Some fluorescence in wounded cells, none in unwounded cells. Usable in fixed cells. Not very bright. |
| DiBAC4(3) | Membrane impermeable in polarized cell. Binds to proteins within the cell. | Blue / green | Can not be used in fixed cells (bright in both wounded and unwounded cells). In unfixed cells, slowly infiltrates unwounded cells at variable rates between cells. |
| Fluorescein | Membrane impermeable. Biologically inert. | Blue / green | Poor signal. Requires washing steps. |
